# Science Education for the Youth (SEFTY): A Neuroscience Outreach Program for High School Students in Southern Nevada During the COVID-19 Pandemic

**DOI:** 10.1101/2024.02.02.578716

**Authors:** Nabih Ghani, Hayley Baker, Audrey Huntsinger, Tiffany Chen, Tiffany D Familara, Jose Yani Itorralba, Fritz Vanderford, Xiaowei Zhuang, Ching-Lan Chang, Van Vo, Edwin C. Oh

## Abstract

Laboratory outreach programs for K-12 students in the United States from 2020-2022 were suspended or delayed due to COVID-19 restrictions. While Southern Nevada also observed similar closures for onsite programs, we and others hypothesized that in-person laboratory activities could be prioritized after increasing vaccine doses were available to the public and masking was encouraged. Here, we describe how the Laboratory of Neurogenetics and Precision Medicine at the University of Nevada Las Vegas (UNLV) collaborated with administrators from a local school district to conduct training activities for high school students during the COVID-19 pandemic. The Science Education for the Youth (SEFTY) program’s curriculum was constructed to incorporate experiential learning, fostering collaboration and peer-to-peer knowledge exchange. Leveraging neuroscience tools from our UNLV laboratory, we engaged with 117 high school applicants from 2021-2022. Our recruitment efforts yielded a diverse cohort, with >41% Pacific Islander and Asian students, >9% African American students, and >12% multiracial students. We assessed the impact of the SEFTY program through pre- and post-assessment student evaluations, revealing a significant improvement of 20.3% in science proficiency (*p*<0.001) after participating in the program. Collectively, our laboratory curriculum offers valuable insights into the capacity of an outreach program to actively foster diversity and cultivate opportunities for academic excellence, even in the challenging context of a global pandemic.

**Significance Statement:** The Science Education for the Youth (SEFTY) program at UNLV successfully engaged 117 diverse high school students in neuroscience-based experiential learning, demonstrating the viability of in-person education during a pandemic. Significant improvements in science proficiency (20.3% increase) underscore the program’s effectiveness in fostering academic excellence and diversity. This initiative potentially serves as a model for maintaining high-quality, inclusive science education in challenging times.

## Introduction

The COVID-19 pandemic transformed the education landscape in the United States, creating potential gaps in knowledge acquisition. In-person instruction and activities faced closures, and access to after-school and extracurricular programs remained limited. As classes were forced online, students struggled to receive the same quality of education and social interaction as they did in person.^1–5^ Students from communities with lower socioeconomic status were impacted disproportionately by the transition online, often having decreased access to essential resources provided through in-person schooling.^2–4,6–9^ The loss of time spent through in-person education has been described as “unfinished learning,” evidenced by significantly decreased reading and math attainment and higher rates of absenteeism during the 2020-21 school year.^3^ Trends of “unfinished learning” have been observed across the United States, with the loss of in-person learning affecting educational opportunities as communities were required to take measures in response to the COVID-19 pandemic. ^1,3,8,9^

Changes to online learning potentially exacerbated local education challenges that were already present. As a result of the pandemic, and similar to what had been seen across the country, reversals in positive trends in education, such as graduation rates, were observed in Nevada. While positive trends are now prevalent in 2023, public education in Nevada has faced challenges in creating opportunities and allocating resources for academic growth that match national averages.^10,11^ This is highlighted partly by the lower teacher-to-student ratio of 1:19 when compared to the national average of 1:16 in 2022.^12^ Additionally, the total amount spent per pupil in the 2020 fiscal year for Nevada was $9,814 compared to a national average of $13,494, although this funding level has trended upward and per pupil expenditures were $10,178 for the 2021-22 school year.^13,14^ The largest of Nevada’s school districts is the Clark County School District (CCSD) in Southern Nevada, which serves over 300,000 students and is the fifth largest school district in the country.^14,15^ CCSD also supports a diverse student population that reflects the general population in Nevada. The school district includes a large geographic area, such as the Las Vegas metropolitan area and surrounding areas in Southern Nevada.^14^ While Southern Nevada is especially diverse, creating programs to promote diversity in STEM fields has required creativity.^10–12,16^

The recruitment of students into STEM fields can be influenced by the size and diversity of school districts. In response to similar challenges across the United States, outreach interventions have been conducted to connect students with STEM and higher education before an undergraduate education.^10,17–19^ Virtual methods, like remote-controlled internet-connected microscopes, can connect students to apparatus that would not regularly be available in school laboratories, allowing them to make real-time observations and draw conclusions based on those observations.^20–22^ With experiential learning, students can engage actively with the ideas they learn conceptually in classrooms or through traditional learning materials. Experiential learning involves participants processing knowledge, skills, and/or attitudes in a learning situation characterized by a high level of active involvement.^23^ A program that utilizes experiential learning actively involves students, allowing them to learn about concepts and apply them in an appropriate environment.^23–26^ One example of such a program is the EvolvingSTEM program, where students actively used laboratory techniques to induce experimental evolution in the bacteria *Pseudomonas fluorescens* and learn about evolution and natural selection. Using experiential learning to examine natural selection promoted a greater understanding of these concepts than students who learned through a standard curriculum using books and lectures.^27^ The NIDDK High School Short-Term Research Experience for Underrepresented Persons is another example of a program where students were involved in research projects to learn more about diverse STEM topics.^28^ Additionally, the Science Education Partnership Award (SEPA) program, supported by the NIH, allows educational organizations to apply for project funding that involves K-12 students in STEM fields. For example, the Maryland Action for Drug Discovery and Pharmaceutical Research (MADDPR) Program is a successful SEPA program that provides lab experience and mentoring for underserved high school students.^29,30^

Local universities have the resources to engage communities and provide valuable STEM education for high school students, opportunities that were lacking due to the COVID-19 pandemic^1,3,8,31–33^. In addition, with increasing access to vaccines and the choice to employ masks, laboratory outreach programs can be conducted responsibly. Here, we describe a program called Science Education for the Youth (SEFTY) to engage high school students to facilitate growth and interest in science. With the SEFTY program, we aimed to create a curriculum incorporating experiential learning to introduce students to common biological concepts while improving their scientific literacy. This was done by combining lectures, laboratory work, and journal club, where students analyzed scientific literature and presented findings from a published manuscript. In the curriculum context, we also worked to facilitate collaboration and team building with students. Finally, we sought to quantify our success in engaging students and improving their scientific understanding.

## Materials and Methods

### Ethics statement

The University of Nevada Las Vegas (UNLV) Institutional Review Board (IRB) reviewed this project and determined it to be exempt from human subject research according to federal regulations and university policy.

### Structure of the Program

The SEFTY program was created to provide high school students access to education relating to biomedical sciences and a hands-on experience with experiments to reinforce their learning. The program began in October 2021 and focused on recruiting 11th and 12th-grade students. All instructors were administered the COVID-19 vaccine and masked during the outreach experience. During the 2021-2022 school year, the SEFTY program ran for eight-week sessions, two days a week. Three sessions were held during the school year, one during the fall semester, one during the spring semester, and a two-week cohort was also conducted during the summer. Another session was conducted in the Fall semester of 2022. The program structure involved a lecture on a particular biomedical topic followed by students conducting an experiment in the lab that demonstrated the topic. Concurrently, students also read and analyzed a scientific paper centered around neurodevelopment. To ensure equitable access for all students, the Nevada INBRE procured twelve Dell Chromebooks 3100, specifically for this educational activity. A road map (**Figure 1**) was shared with students at the beginning of the program that outlined which topics would be covered during the program and how the ideas were related.

**Figure 1.**
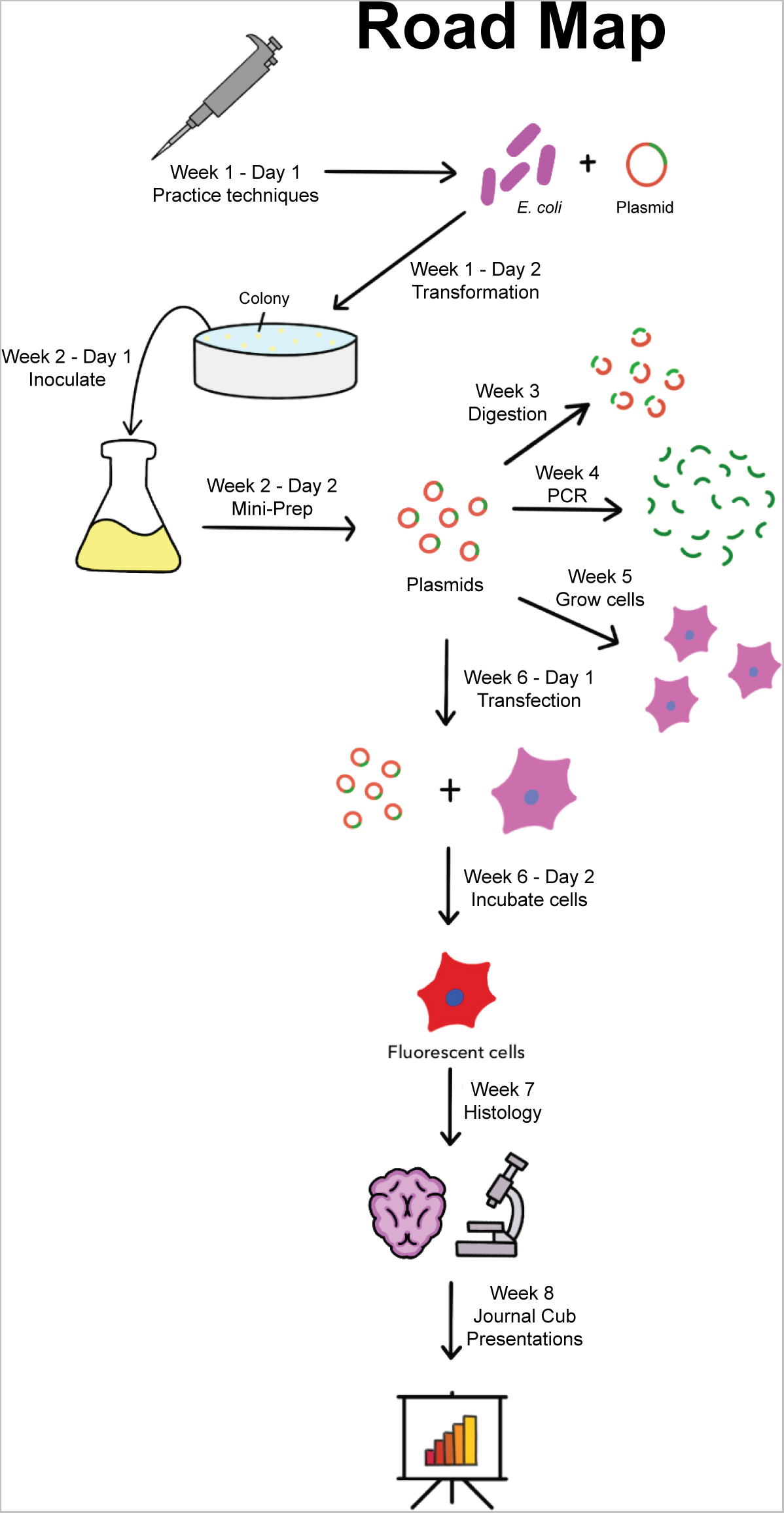
The SEFTY Plan. The roadmap given to students served as a comprehensive overview of the program, contextualizing the various processes they learned. It was crucial to connect the specific skills acquired in the program to broader biological concepts, enhancing their understanding and application.

The program began with students exposed to an introductory activity, a pre-survey, and a pre-assessment. Then, students received a lecture on lab safety, scientific notation, the metric system, and micropipettes and worked together to complete activities in lab manuals to reinforce their learning. After completing these activities, students were oriented to the lab to familiarize themselves with critical lab safety locations and to learn pipetting techniques. Micropipettes (**Figure 2A**) are essential tools in many biology labs and allowed for the demonstration of the metric system and scientific notation they learned. On the first day, students were also introduced to the journal club part of the program, where they received a scientific article to read and present at the end of the program. The following day, students were introduced to the central dogma of biology, plasmids, and aseptic technique.

**Figure 2.**
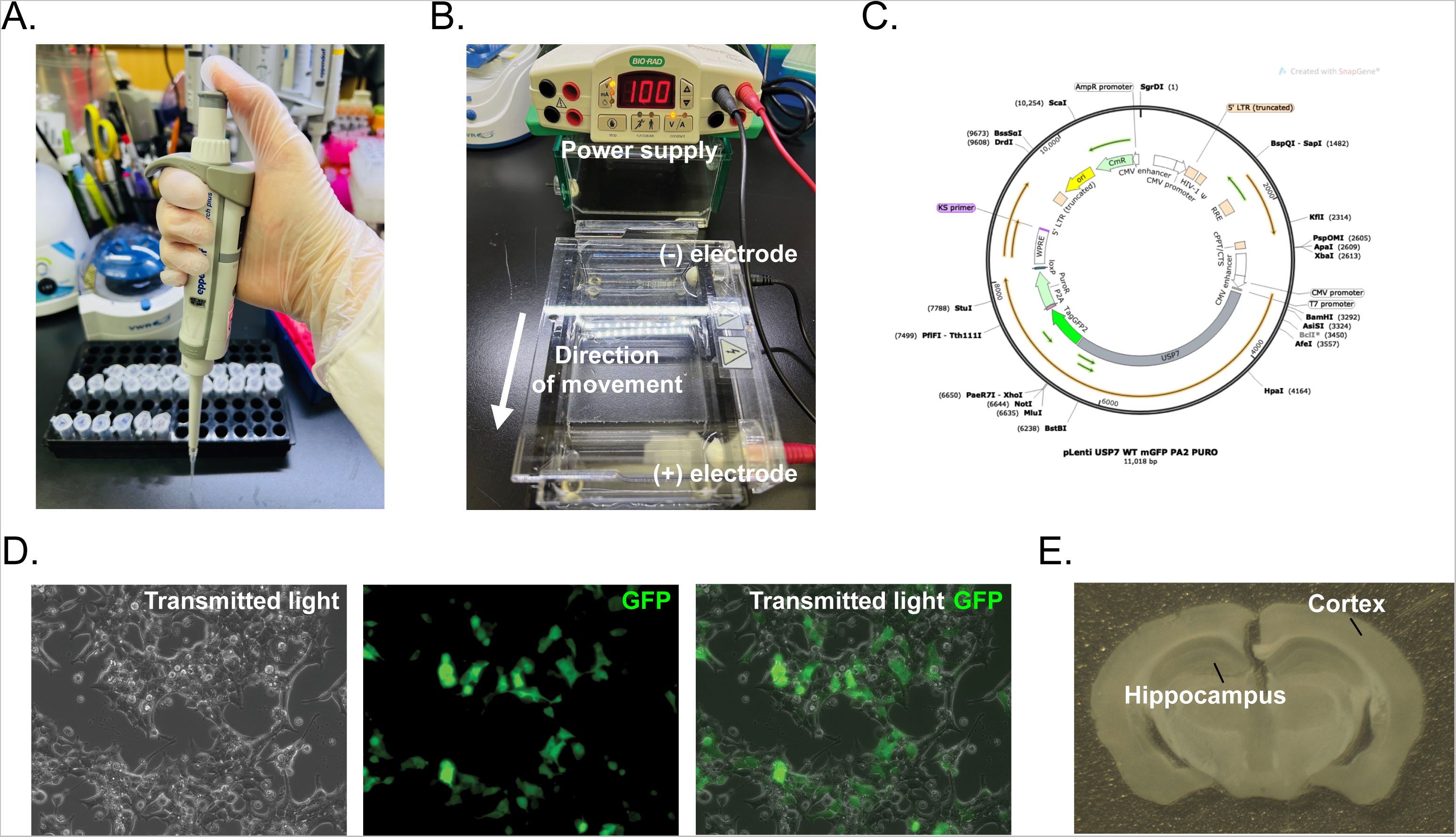
An overview of the technical concepts covered during the SEFTY sessions. (A) Image of a hand demonstrating proper pipetting technique. (B) A figure showing the general principles behind gel electrophoresis, a process that was used to analyze fragments of DNA formed from the digestion of a plasmid. (C) A diagram from SnapGene showing the structure of a plasmid and restriction sites that can be cut for analysis. (D) An image of HEK293T cells under transmitted light microscopy, showing green fluorescent protein (GFP) expression from a plasmid transfected by students. The panel also includes a merged view, combining both the transmitted light and GFP fluorescence. (E) An image of a mouse brain section that was shown to further understanding of the brain’s structure (hippocampus and cortex) and how immunohistochemistry experiments can play a role in understanding biology.

Students then applied these topics to transform bacterial cells with plasmids containing the gene studied in their respective journal club articles. Bacterial colonies were grown on ampicillin plates overnight. The next day, students reviewed the importance of sterile technique and inoculated liquid cultures from the plates they had previously transformed. After allowing their inoculated cultures to grow, students reviewed plasmids and their significance for bacteria in their natural environment. They isolated their respective plasmids that were then used to perform digestion, polymerase chain reaction (PCR), and transfection into HEK293T (human embryonic kidney cells expressing the simian virus 40 (SV40) large T antigen) cells. The papers that were read as a part of the journal club portion of SEFTY used plasmid constructs to examine expression in neurodevelopmental disorders, and the isolated plasmids became essential to most of the remaining experiments conducted in the lab.

Students then learned the basics of plasmid digestion by restriction enzymes and gel electrophoresis. They were given a map of the plasmid and were asked to predict what bands would appear in an agarose gel based on the restriction enzymes used (**Figure 2B-C**). This also allowed students to understand further the structure of DNA in prokaryotes versus eukaryotes and the chemical structure of DNA that allows for its migration in agarose gels. The following week, students learned about PCR. They used the plasmids they isolated to conduct a reaction to amplify DNA and observed the amplification of the gene of interest through gel electrophoresis.

Following digestion and PCR, students were taught about cell culture and its utility within a laboratory. As HEK293T cells were used for cell culture experiments, students were able to review the differences between eukaryotes and prokaryotes. Through cell culture, students learned about why cell lines are used frequently in laboratories and the advantages that they confer. Additionally, they learned about the origins of cell culture techniques, such as the HeLa cell line, and discussed ethical considerations in science. Students then transfected their cells with their plasmid, allowing for the overexpression of their gene of interest, which had a fluorescent tag of green fluorescent protein (GFP) (**Figure 2D**). They observed the fluorescence of the cells that were transfected, and through this, they were also able to learn and apply the basics of handling microscopes in a laboratory.

As their final experience in the lab, students learned about histology and the functions of the different lobes in the brain. They could look at mouse brain sections to identify structures like the hippocampus and cortex (**Figure 2E**). Understanding the brain’s structure was an important point covered in the journal club articles that focused on neurodevelopment and analyzed changes in brain anatomy. Towards the end of the SEFTY program, students presented the paper they had been reading. These laboratory experiments allowed students to take advantage of experiential learning and apply knowledge to reinforce what they learned in the program.

### Lesson Plan

Complete work modules and the SEFTY lesson plan (**Supplementary Data 1**) were prepared before each new SEFTY session for the semester.

### Evaluation Plan and Statistics

A Google and paper form were used to survey science proficiency before and after each SEFTY cohort. The pre-evaluation and post-evaluation survey contained multiple-choice questions (**Supplementary Data 2**) to test knowledge and the impact of the SEFTY outreach program. The percentage of multiple-choice questions answered correctly in the pre and post-tests were computed. For students enrolled in each session, a paired t-test was used to assess the pre- and post-performance differences. To evaluate the overall SEFTY program effect across all sessions, the improvement percentage was computed for each student. A linear regression model was then used to test if there was a significant improvement accounting for the sessions. Graphs were generated in GraphPad Prism.

## Results

### Recruitment outcomes

Diversity in STEM can help promote innovation and fairness and minimize inequities.^11,34^ This is especially true in a changing American landscape that is projected to be composed of an increased number of people identifying with multiple racial groups.^35^ Southern Nevada especially boasts an increasingly diverse student population (**Figure 3A**). To assess our outreach to diverse student populations in our community, we gathered data about applicants (n=117) and students (n=34) in our program. Our analysis revealed 11.97% of applicants and 11.76% of students identified as multiracial compared to 7.55% of students in all of the local school district (**Figure 3A-C**).^14^ Furthermore, while Hispanic students comprise the largest percentage of students in the district at 47.7%, only 9.40% of applicants and 2.94% of students engaged by our program were Hispanic (**Figure 3A-C**).^14^ Pacific Islanders and Asians were overrepresented in both applicants and students of the program (**Figure 3B-C**). The results of these analyses identify recruitment as a potential target for improvement of our outreach program.

**Figure 3.**
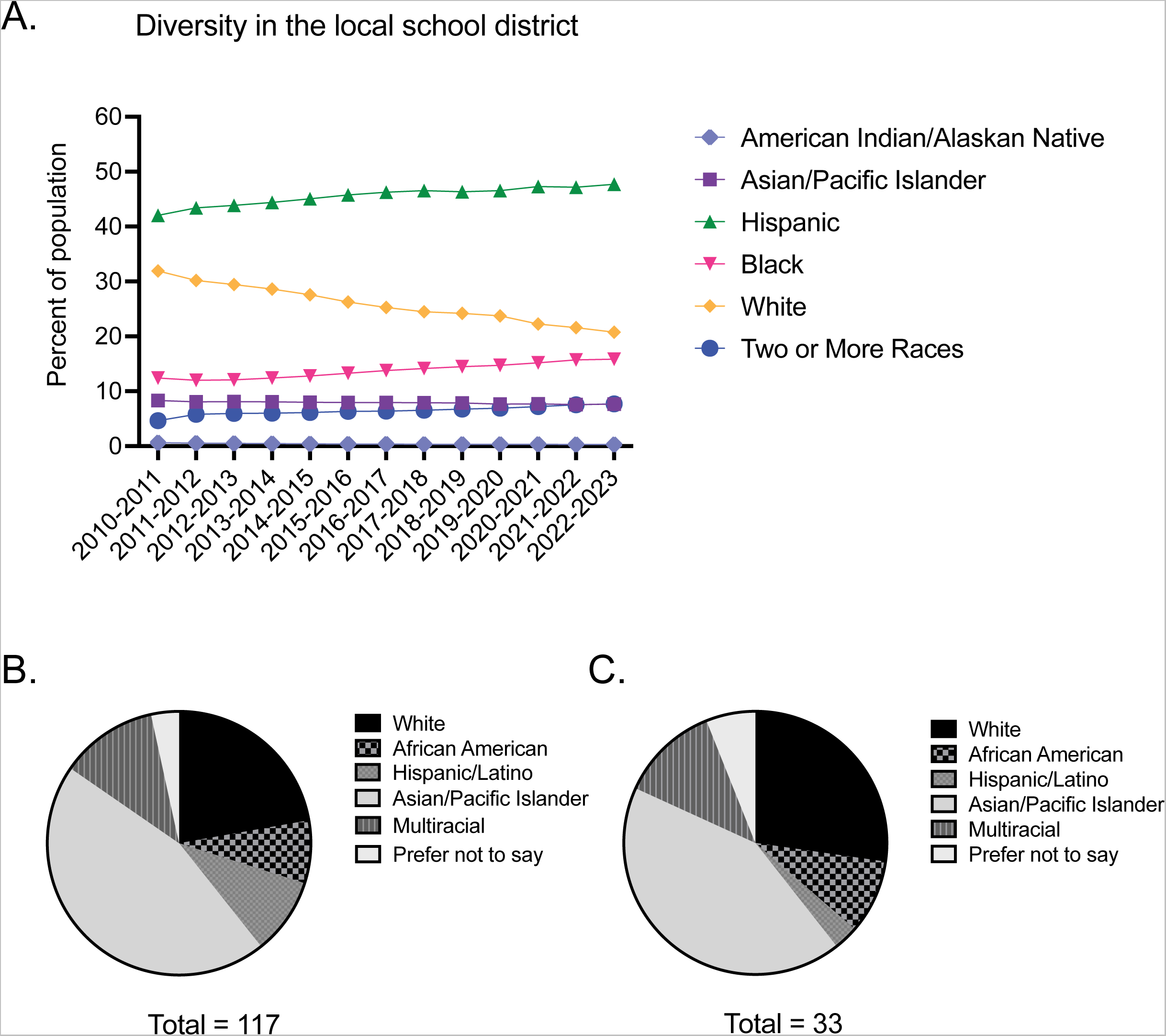
Diversity in K-12 school district and SEFTY. (A) This graph tracks the changing diversity based on racial background within CCSD since the 2010-2011 school year. (B) This chart displays the racial demographic data of the applicants to the SEFTY program. (C) This chart shows the racial demographic data for the students who were accepted and participated in the SEFTY program.

Students applied for the program through a form submitted to multiple schools in the community. The students were selected based on a short personal statement about their interest in the program. Students were also only accepted if they were in 11^th^ or 12^th^ grade and if they had transportation to the program. As over 120,000 students in the local school district take school buses, transportation was a limiting factor in the ability of students to attend the SEFTY Program.^36^

### Using peer learning to strengthen scientific literacy and soft skills

During their participation in the program, students focused on enhancing their scientific literacy by studying a research paper related to the gene central to their lab experiments. To ensure equal access to information, Dell Chromebooks with UNLV-provided Wi-Fi access were made available to all students. The program culminated with each student delivering a presentation, summarizing their insights from both the paper and their overall learning experience. To prepare students for their presentations, they worked together to read the paper with the help of the program leaders. Additionally, peer learning was implemented on different topics throughout the program. In this paper, peer learning is defined as students teaching and learning from one another without direct instruction from a program leader or teacher.^37,38^ Students would act as both students and teachers, studying a specific topic and presenting it to their peers. Our goal with the peer learning exercises was to create a collaborative environment and allow them to practice communicating ideas related to STEM. Specifically, this approach was used to explore concepts such as the differences between DNA and RNA and the central dogma of biology. A group of two or three students would provide explanations and present what they independently learned to the rest of the students. This was also done to help students understand the differences between different neurological disorders and how they may exhibit phenotypes in patients with these genetic disorders. With this foundational work of collaboration and presentations, students worked together to break down a paper that utilized techniques they encountered in the lab and present it to an audience as the program’s culmination.

Students from eight high schools in Southern Nevada participated in the program. Our final goal of the program was a group presentation on a paper they were assigned, so facilitating the collaboration for the students was essential to ensure success and growth. Collaborative learning efforts, such as peer learning, can trigger critical thinking and are related to students’ motivation.^39^ Additionally, students’ ability to interact with one another was limited due to the COVID-19 pandemic, which has been commonly cited as a significant disadvantage of virtual education during the pandemic.^1,5,6,8^ One way we aimed to create an environment where students were comfortable discussing and working with one another was through a low student-to-program leader ratio. It has been well established that a lower student-to-teacher ratio and the interactions between students and teachers, and in this case program leaders, allow for a better experience in a class setting and improved attitudes and performance of students.^40–42^ For most of the program, one or two program leaders were responsible for the cohort, typically seven or eight students. Integrating discussions, asking questions of the students, and having them present, as discussed previously, were mechanisms used frequently to help build comfort and subsequent collaboration in an unfamiliar environment. Finally, activities like group scavenger hunts on the University of Nevada, Las Vegas campus, and photo contests were undertaken to allow students to feel comfortable in front of each other, foster a collaborative environment, and enjoy their learning experience as a part of the SEFTY program.

### Increasing understanding of science through the SEFTY Program

As the program’s primary goal is to educate students, an assessment was designed to understand growth through the program. The multiple-choice assessment was administered on the program’s first and final day (see **materials and methods**). Following the end of the program, the scores were compared to determine whether students improved as a result of the program. Significant improvement on test performances were observed for Fall 2021 (t(df=6)=3.90, *p*=0.008), Spring 2022 (t(df=7)=5.79, *p*<0.001) and Fall 2022 (t(df=6)=2.90, *p*=0.03) sessions (**Figure 4A**). Overall, the post-assessment test showed an average of a 20.3±13.5% increase (t(df=19)=4.52; *p*<0.001) from the pre-assessment test across all sessions. Additionally, students completed surveys to assess the program’s effectiveness and how it impacted their interests in biology. Students were asked to rate their interest in biology before and after the program on a scale of 1-10, 1 being not very interested and 10 being very interested (**Figure 4B**). On average, students who completed the surveys (n=16 in one session) rated their interest in biology a value of 7.69±2.02, which increased to an average rating of interest of 8.94±1.24 at the end of the program (*p*=0.001) (**Figure 4B**).

**Figure 4.**
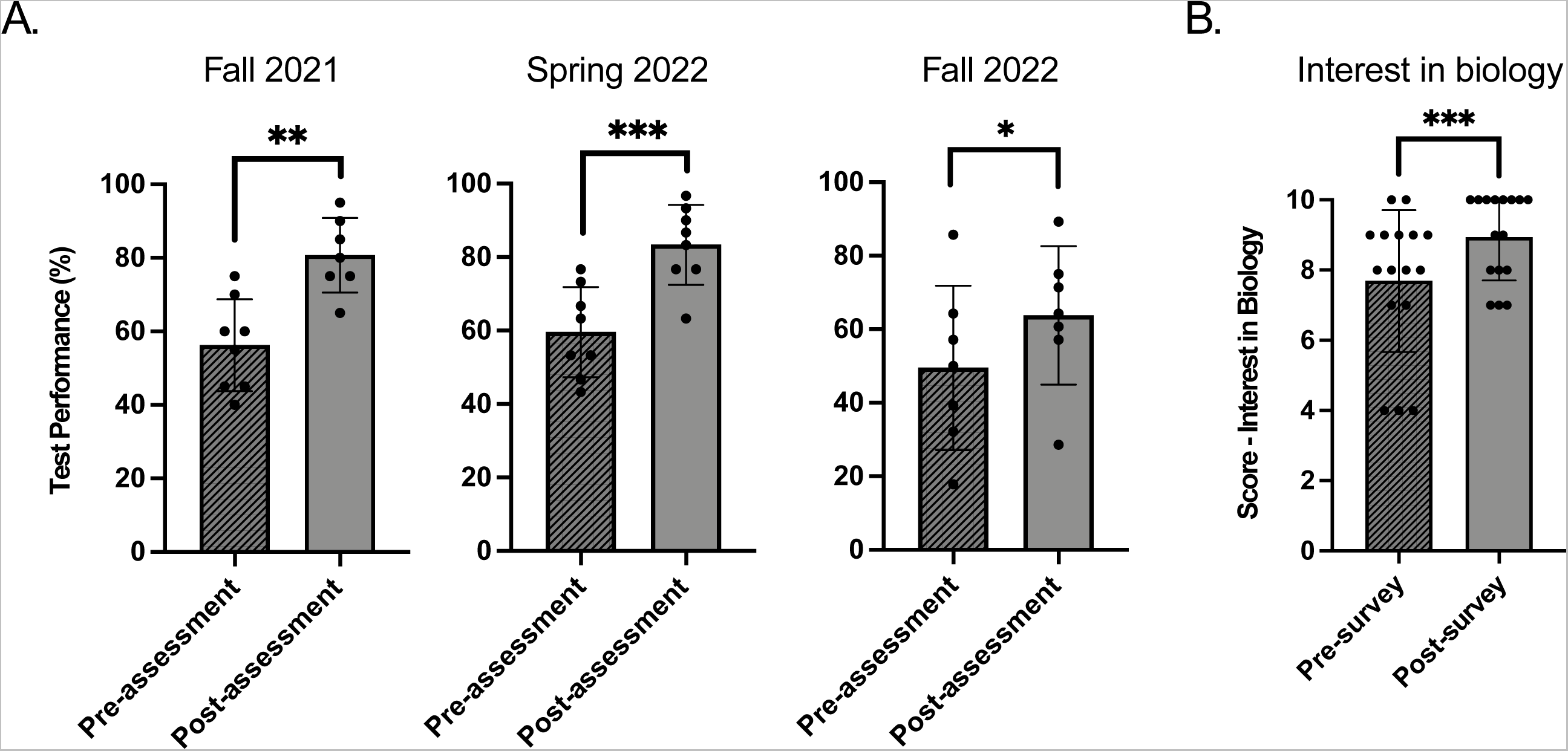
Assessment of the SEFTY impact. (A) This figure shows the difference between the averages for the pre and post-assessments for students in all three cohorts. (B) For the 16 students who completed both pre and post-program surveys, this figure shows the average response to the survey question: “On a scale of 1-10, how interested are you in biology?” An asterisk (*) refers to *p*≤0.05, two asterisks (**) refers to *p*≤0.01, and three asterisks (**) refers to *p*≤0.001.

## Discussion

The execution of the SEFTY Program during the COVID-19 pandemic revealed critical insights into the challenges and opportunities in recruiting a diverse student body. While the program successfully engaged a significant proportion of multiracial students, exceeding the school district average, it underperformed in attracting Hispanic students, who form the largest demographic in local schools (**Figure 3**). This disparity highlights a pressing need to refine recruitment strategies. The underrepresentation of Hispanic students, despite their prevalence in the community, suggests that current outreach efforts may not effectively resonate with this demographic. To bridge this gap, a multifaceted approach is essential. Firstly, enhancing collaboration with local schools and community organizations can provide more targeted and culturally sensitive outreach.^31,43,44^ This involves not only advertising the program more effectively but also understanding and addressing the unique challenges faced by these students, such as transportation issues. Additionally, seeking recommendations from teachers could uncover hidden talents and interests, especially in students who might not typically apply due to perceived barriers.^16,43^ Implementing such strategies could significantly enhance the diversity of the program, aligning it more closely with the community’s demographic makeup and enriching the educational experience for all participants.

### Enhancing Learning through Collaborative and Peer-Led Approaches

The program’s emphasis on peer learning and collaborative efforts in scientific literacy and soft skills development marks a step towards a more inclusive and effective educational model. This approach not only accommodates diverse learning styles but also fosters a sense of community and shared purpose among the students. The success of this method is evident in the students’ improved ability to understand and articulate complex scientific concepts, as demonstrated in their group presentations and enhanced test scores. However, the constraints imposed by the COVID-19 pandemic have underscored the need for adaptive and resilient educational strategies.^1,3,8,21,22,29^ In future iterations of the program, blending online and in-person learning modalities could offer a more flexible and accessible platform for students, overcoming physical and geographical barriers.^21,22,29^ Additionally, maintaining a low student-to-program leader ratio is crucial in providing personalized support and feedback, which is particularly beneficial in a diverse learning environment.^40–42,45^ These adaptations can ensure that the program remains effective and inclusive, regardless of external challenges, thereby continuing to foster a supportive and collaborative learning atmosphere.

### Demonstrating Impact and Fostering Future Growth in STEM

The significant improvement in students’ test scores and their increased interest in biology as a result of their participation in the SEFTY program highlights its effectiveness in enhancing scientific understanding and fostering a passion for STEM fields (**Figure 4**). This positive outcome may be in response to the program’s well-structured curriculum and its emphasis on active learning and peer engagement. Looking forward, the program’s success in improving scientific literacy and enthusiasm for biology among its participants can serve as a model for similar initiatives. To build on this success, it would be beneficial to conduct longitudinal studies to track the long-term impact of the program on students’ academic and career trajectories in STEM fields. Such data would provide valuable insights into the lasting effects of the program and help identify areas for further improvement. Interestingly, data from our cohort of SEFTY students indicate that the majority of participants have graduated from high school. Ultimately, by continuing to evolve and adapt to the needs of its participants, the SEFTY Program can play a pivotal role in cultivating the next generation of diverse and talented STEM professionals.

As the integration of neuroscience and genetics becomes more prevalent in health-related treatments, the SEFTY program aims to include genomics—the study of an individual’s complete DNA structure and expression—in its curriculum.^46^ Given the expanding scope of genomics in the biomedical field, it encounters challenges similar to those in other biomedical careers, as highlighted in an NIH draft report on diversity in these fields. ^34,47,48^ By incorporating genomics, the program aims to spark and expand interest in STEM more broadly, leveraging the appeal of this growing field. ^46–49^ Overall, our outreach program demonstrates the effectiveness of laboratory initiatives in promoting student diversity and enhancing scientific literacy among K-12 students.

## Acknowledgments

ECO and VV are supported by NIH grants: GM103440 and MH109706, a CARES Act grant from the Nevada Governor’s Office of Economic Development and Grant Number NH75OT000057-01-00 from the Centers for Disease Control and Prevention. The project contents are solely the responsibility of the authors and do not necessarily represent the official views of the Centers for Disease Control and Prevention. We would like to acknowledge personnel at the local school district who shared information about the SEFTY program with students.

