## Supplemental Data for "Science Education for the Youth (SEFTY): A Neuroscience Outreach Program for High School Students in Southern Nevada During the COVID-19 Pandemic"

**Supplementary Data 1**

| Schedule | | |
|---|---|---|
| Week 1 | Thursday (10/6) | Introduction to the program, supplies, and metric system<br>Practice lab techniques<br>Transformation |
| | Friday (10/7) | Introduction to biology, the central dogma of biology, importance of genetics<br>Choose papers for Journal Club<br>How to approach reading peer-reviewed articles |
| Week 2 | Thursday (10/13) | Sterile technique, inoculation, and media<br>Scavenger hunt around UNLV campus |
| | Friday (10/14) | Miniprep and nanodrop |
| Week 3 | Thrsday (10/20) | Digestion and running digestion products on a gel |
| | Friday (10/21) | Discuss results, repeat experiment if needed |
| Week 4 | Thursday (10/27) | Polymerase Chain Reaction<br>*Gattaca* movie |
| | Friday (10/28) | PCR gel electrophoresis and imaging |
| Week 5 | Thursday (11/3) | Introduction to cell culture (splitting cells) |
| | Friday (11/4) | View cells<br>Genetic disorder presentation<br>Work on Journal Club articles |
| Week 6 | Thursday (11/10) | Transfection<br>*Human Nature* movie |
| | Friday (11/11) | View cells and image<br>Constructing a CV, crafting professional emails, cover sheets |
| Week 7 | Thursday (11/17) | Introduction to histology and brain; view sections/image<br>Match regions to atlas of brain and match regions to disease |
| | Friday (11/18) | Review stations |
| No SEFTY this week - Thanksgiving break | | |
| Week 8 | Thursday (12/1) | Journal Club Practice |

| | | |
|---|---|---|
| | Friday (12/2) | Journal Club Presentations |

**Supplementary Data 2**

Name: ____________________________________________ Date:____________________

### Pre- and post-assessment

1. Convert 11 L to mL
    a. 11,000 mL
    b. 1.1 mL
    c. 110,000 mL
    d. 1100 mL

2. What are the three domains of life?
    a. Eukarya, Bacteria, Archaea
    b. Eukarya, Prokarya, Bacteria
    c. Fungi, Animalia, Plantae
    d. Bacteria, Archaea, Animalia

3. How many microliters are within one liter?
    a. 1 x $10^{-3}$ $\mu L$
    b. 1 x $10^{-2}$ $\mu L$
    c. 1 x $10^{3}$ $\mu L$
    d. 1 x $10^{6}$ $\mu L$

4. Which of the following is an <u>incorrect</u> nucleotide base pairing?
    a. A-G
    b. G-C
    c. A-T
    d. A-U

5. Which of the following nucleotides are not found in DNA?
    a. Adenine
    b. Guanine
    c. Thymine
    d. Uracil

6. Plasmids are composed of which type of biological molecule?
    a. Nucleotides
    b. Proteins
    c. Amino acids
    d. Carbohydrates

7. Which of the following contains all of the required genetic information for a bacterial cell to survive?
    a. Plasmid
    b. Gene
    c. Chromosome
    d. Ribosome

8. By what process do bacteria cells divide?
    a. Mitosis
    b. Meiosis
    c. Binary fission
    d. Sexual Reproduction

9. Where is the DNA kept inside of a bacterial cell?
    a. Nucleus
    b. Nucleolus
    c. Nucleoid Region
    d. Golgi Apparatus

10. What three processes are components of the Central Dogma of Biology?
    a. Transformation, Mitosis, Translation
    b. Replication, Transcription, Translation
    c. Replication, Meiosis, Transcription
    d. Interphase, Mitosis, Cytokinesis

11. In eukaryotic cells, transcription takes place in the _________, while translation takes place at ribosomes found in the __________.
    a. nucleus; cytoplasm
    b. cytoplasm; nucleus
    c. nucleolus; nucleus
    d. cytoplasm; cytosol

12. Which of the following biological molecules help catalyze chemical reactions?
    a. Phospholipids
    b. Channel Proteins
    c. Nucleic Acids
    d. Enzymes

13. What is a gene?
    a. A region of a protein
    b. A region of DNA
    c. All of the DNA found in an organism
    d. A region of RNA

14. How many pairs of chromosomes do humans have?
    a. 46
    b. 23
    c. 50
    d. 25

15. What is a restriction enzyme?
    a. An enzyme isolated from bacteria that cuts DNA at a certain sequence
    b. An enzyme isolated from protists that cuts DNA at a certain sequence
    c. An enzyme that splices together strands of DNA
    d. An enzyme that unwinds DNA strands during DNA replication

16. After 3 rounds of PCR how many molecules of DNA will have formed from an original DNA molecule?
    a. 2
    b. 4
    c. 6
    d. 8

17. What is a difference between chromosomes in eukaryotic cells and prokaryotic cells?
    a. Eukaryotic cells have much smaller chromosomes than prokaryotic cells
    b. Chromosomes in prokaryotic cells are made of RNA, but chromosomes in eukaryotes are composed of DNA
    c. Prokaryotic cells typically have two sets of chromosomes, while eukaryotes have four sets; two from each parent
    d. Chromosomes are found in the nucleus in eukaryotic cells and the cytoplasm in prokaryotic cells

18. How many nucleotides code for a single amino acid during translation?
    a. 1
    b. 2
    c. 3
    d. 6

19. In which step of PCR do the strands separate?
    a. Excision
    b. Denaturation

c. Annealing
d. Extension/elongation

20. What type of enzyme is used to elongate DNA strands during PCR?
    a. RNA polymerase
    b. DNA Polymerase
    c. Helicase
    d. Isomerase

21. Which of the following is **NOT** characteristic of bacteria?
    a. Membrane bound organelles
    b. Unicellular organisms
    c. Circular DNA
    d. Lacking a nucleus

22. How many lobes is the human brain made of?
    a. 2
    b. 3
    c. 4
    d. 5

23. In which of the following parts of the brain does most information processing, consolidation, and cognition occur?
    a. Cerebrum
    b. Cerebellum
    c. Brain Stem
    d. Midbrain

24. Which of the following parts of the brain is most closely paired with its associated function?
    a. Cerebellum; cognitive functions and planning
    b. Occipital lobe; processing visual information
    c. Pyriform lobe; coordinates movement
    d. Temporal lobe; cognitive functions and planning

25. Which of the following is **NOT** a reason for using mouse models instead of humans?
    a. Mice are less expensive
    b. Mice and humans have similar physiological and structural qualities
    c. Mice allow for a better understanding of complete genetic profile in comparison to other commonly used animals in research
    d. Mice are biologically equivalent to humans as embryos

26. During gel electrophoresis of DNA, DNA in the gel travels from the ___________ anode towards the ___________ cathode.

a. larger; smaller
b. smaller; larger
c. negative; positive
d. positive; negative

27. What chemical property of DNA allows for it to migrate in a gel via gel electrophoresis?
    a. Its chemical interactions with agarose
    b. The pairing of the nitrogenous bases
    c. The positive charge of the ribose sugars in the sugar-phosphate backbone
    d. The negative charge of the phosphate groups in the sugar-phosphate backbone

28. Scientist Jerry is setting up a digestion of a plasmid. He wants to use 1000 ng of his plasmid, which has a concentration of 400 ng/$\mu L$. What volume of plasmid should he use to set up his reaction?
    a. 2.5 $\mu$g
    b. 25 $\mu$g
    c. 25 $\mu$L
    d. 2.5 $\mu$L

Graph the following data. Be sure to include axis labels, a title, and a legend.

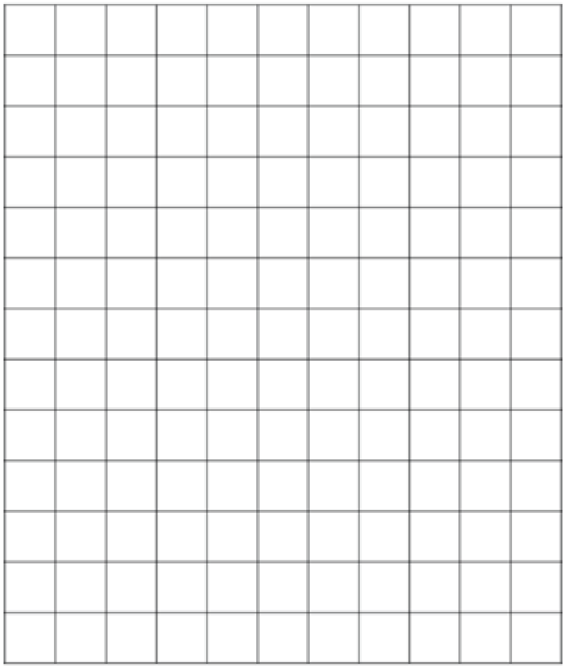

| | GPA | |
|---|---|---|
| **Hours Spent Studying** | **Student 1** | **Student 2** |
| **2** | 1.5 | 1.0 |
| **4** | 2.5 | 2.5 |
| **6** | 3.5 | 3.0 |
| **8** | 4.0 | 4.0 |

Widow’s peaks are a dominant trait in people. A heterozygous male with a widow’s peak reproduced with a homozygous recessive female. What are the possible genotype and phenotype percentages for their offspring?

Genotypes:

AA: _______ %

Aa: _______ %

aa: _______ %

Phenotypes:

Widow’s peak: _______ %

No widow’s peak: _______ %
